# Conscious and Unconscious Emotional Processing: Insights from Behavioral and Pupillary Responses

**DOI:** 10.1101/2024.07.20.604402

**Authors:** Helia Taghavi, Leila Sharbaf, Fatemeh Ismailzad, Abdol-Hossein Vahabie, Javad Hatami, Ehsan Rezayat

## Abstract

Emotions are crucial in social interactions, influencing communication and relationships. Distinguishing the perceived emotion in conscious and unconscious emotional processing is a key research area with cognitive and physiological implications. This study investigates conscious and unconscious emotional processing through behavioral and pupillary responses. Participants completed emotion recognition tasks under varying states, revealing higher accuracy in conscious emotion identification. Emotions like anger, happiness, fear, surprise, and neutral elicited distinct response patterns. Pupillometry data showed pupil size suppression in the conscious state and enhancement in the unconscious state, with differences in peak pupil size across emotions. Task-related components, amplitude, and latency parameters differed between conscious and unconscious states, highlighting the role of awareness in emotional regulation. These findings emphasize the complex interplay of cognitive and physiological processes in emotional responses, providing insights into emotional recognition mechanisms. This study contributes to understanding emotional processing dynamics and has implications for psychology and neuroscience research.

## Introduction

Emotions play a crucial role in human social interactions, serving as a means of fast and effective communication. Accurately perceiving emotional cues conveyed through facial expressions, tone of voice, and body language is vital for social competence and fostering positive relationships (Bornhofen & Mcdonald, 2008). Understanding facial affect is fundamental to comprehending others’ emotions and navigating social interactions (Spikman et al., 2013).

Emotional processing encompasses both conscious and unconscious aspects, each associated with distinct neural responses (Kihlstrom et al., 2000). Unconscious emotions manifest in two forms: those generated without conscious awareness of their source and implicit emotions that occur without entering consciousness (Smith & Lane, 2016).

Neuroscientific studies have found notable differences in neural responses during the processing of conscious and unconscious emotions. Functional magnetic resonance imaging (fMRI) research indicates activation of distinct brain regions for each type of processing, illuminating the involvement of different neural pathways (Phillips et al., 2004; Tamietto et al., 2015). Moreover, event-related potentials (ERPs) reveal unique temporal patterns in neural responses to conscious and unconscious perception of emotions, underscoring the qualitative disparities in processing (Liddell et al., 2004; Williams et al., 2004). In addition to neuroimaging approaches, physiological studies employing measures like electromyography (EMG) shed light on the effects of conscious and unconscious emotional processing. Such studies have supported the notion that conscious perception exerts inhibitory influence on subcortical pathways, providing insights into the interplay between cortical and subcortical processing (Dimberg et al., 2000; Tamietto et al., 2009).

The relationship between emotions and physiological arousal has been elucidated by the James-Lange theory, emphasizing the role of bodily feedback in shaping emotional experiences (James, 1948; Tamietto et al., 2015). Pupillometry, which measures changes in pupil size, has emerged as a valuable tool in investigating conscious and unconscious emotional processing, offering insights into cognitive load, attention, and emotional processes (Bradley et al., 2008). Studies investigating recognition thresholds for facial emotions across different age groups have unveiled variations in the ability to recognize both conscious and unconscious emotions, highlighting the impact of age on emotional perception (Calder et al., 2003; Laeng et al., 2012). Given the gaps in understanding the impact of different emotional expressions on pupil size, this research seeks to explore how consciously and unconsciously perceived emotions influence pupil size. The study further aims to investigate whether presenting various emotions consciously versus unconsciously results in distinct effects on pupil size and whether specific patterns in pupil size can be observed during the unconscious presentation of the six basic emotions.

## Method

### Participants

A total of 30 healthy participants (mean age=26.32, SD=5.68, range 20-39) were recruited for this study. Prior to the experiment, all participants underwent vision screening and completed the Spielberger state-trait questionnaire. Written informed consent was obtained from all participants, and they were compensated upon completion of the experiment. Two participants were excluded from the analysis due to incomplete data. The study was approved by the Ethical Committee of the Iran University of Medical Sciences.

### Stimuli and Apparatus

Emotional and neutral facial expressions were sourced from the Warsaw data set (Olszanowski et al., 2015). To ensure consistency in luminance across stimuli, image editing was performed using Photoshop software. Backgrounds and distracting elements in the photos were removed, and a standardized gray background was applied. Luminance and histogram matching were conducted using the SHINE toolbox (Willenbockel et al., 2010) within MATLAB. Twenty identities (10 male) were selected as stimuli for the behavioral task, and emotional state names were prepared using Photoshop. All experimental tasks were programmed using PsychToolbox-3 in MATLAB. Eye signals were recorded and processed using the EyeLink 1000 eye tracker, positioned 55 cm from the participant’s chin rest with a monitor-to-participant distance of 60 cm.

### The computerized task of perceiving facial emotion

Participants engaged in a facial emotion perception task that presented 14 emotional states derived from male and female identities on a monitor screen. The task involved two stages - conscious facial expression perception and unconscious facial expression perception. Prior to the main experiment, participants underwent training trials to familiarize themselves with the task.

#### Conscious facial expression perception

This task focused on investigating conscious perception of facial emotions and its impact on pupil size. Emotional expressions were presented randomly, followed by a response page where participants selected the corresponding emotion name. The trial sequence involved a 1000 ms fixation cross, a 1000 ms emotional face stimulus, a 500 ms delay, and the response selection. Participants completed 210 trials, each repeated 15 times, lasting approximately 16-20 minutes.

#### Unconscious facial expression perception

In this task, emotional faces were briefly presented to explore the effect of unconscious emotion perception on pupil size. A neutral mask immediately followed to prevent conscious perception. The trial sequence consisted of a 1000 ms fixation cross, a 50 ms emotional face presentation, a 950 ms neutral mask, and the participant’s emotion selection. This task comprised 210 trials, following a similar structure to the conscious perception task, lasting approximately 20 minutes.

### General linear modeling of pupil response

Pupil responses were analyzed using models with 6/8 parameters for conscious and unconscious conditions. The Pupil Response Estimation Toolbox (PRET) (Denison et al., 2020) was utilized for parameter estimation, including internal signal amplitudes and latencies, task-related response amplitudes, linear drift parameters, and baseline shifts. Each event-related signal was characterized by amplitude and latency parameters, reflecting the strength and timing of the internal signal and associated pupil response component, respectively.

### Statistical analysis

Nonparametric tests, specifically the Wilcoxon rank test for pairwise comparisons and ANOVA for overall analysis, were employed to analyze the data.

## Result

### Behavioral Results

Participants’ performance in both conscious and unconscious states was assessed (Figure 2), showing superior performance in the conscious state (p < 0.05), with a score exceeding 70%. Significant differences among emotions were observed in the conscious state (p < 0.01), highlighting disparities between anger and happiness, fear and happiness, as well as neutral and surprise (p < 0.05). The unconscious state analysis (p < 0.01) revealed notable differences, particularly between happiness and other emotions except for neutral and surprise. Additionally, distinctions were observed between neutral and all emotions except happiness and surprise, with surprise significantly differing from anger and sadness (p < 0.05).

**FIGURE 1.**
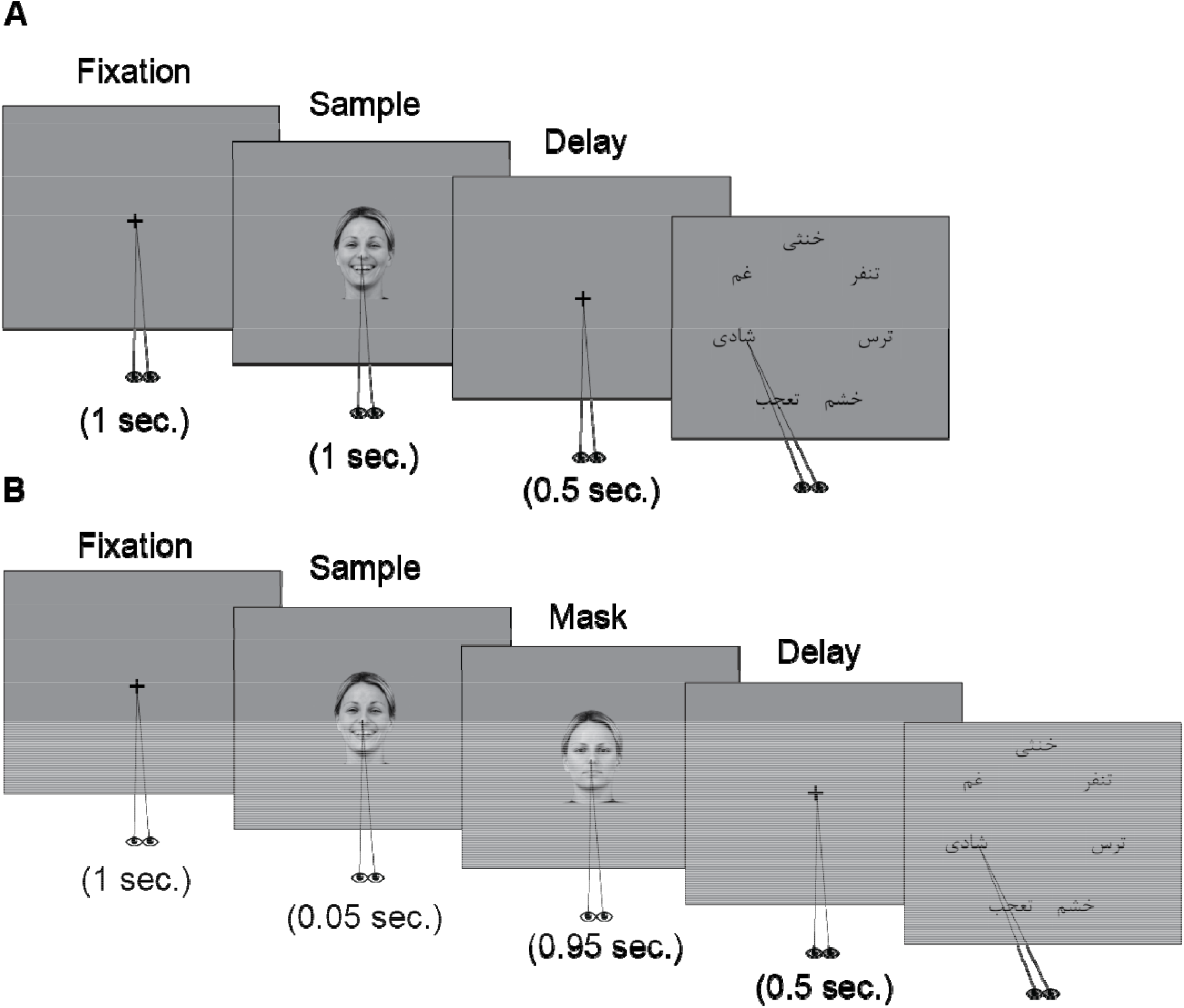
The schematic description of the trial sequences for behavioural tasks. **(A)** For conscious facial expression perception blocks, each trial began with a 1000 ms fixation cross, followed by a 1000 ms presentation of a facial expression, and then a 500 ms fixation cross. Then a response page appeared, and participants responded by saccadic eye movement to select the desired emotion name (= neutral, = sadness, = happiness, = surprise, = anger, = fear, = disgust) and pushed the response key on the keyboard to finalize the response. **(B)** In unconscious facial expression perception trials, a 1000 ms fixation cross preceded a facial expression presented for only 50 ms, which was then masked by a neutral image of the same face for 950 ms, followed by a 500 ms fixation cross. Then the response page appeared until the participant made a response.

**FIGURE 2.**
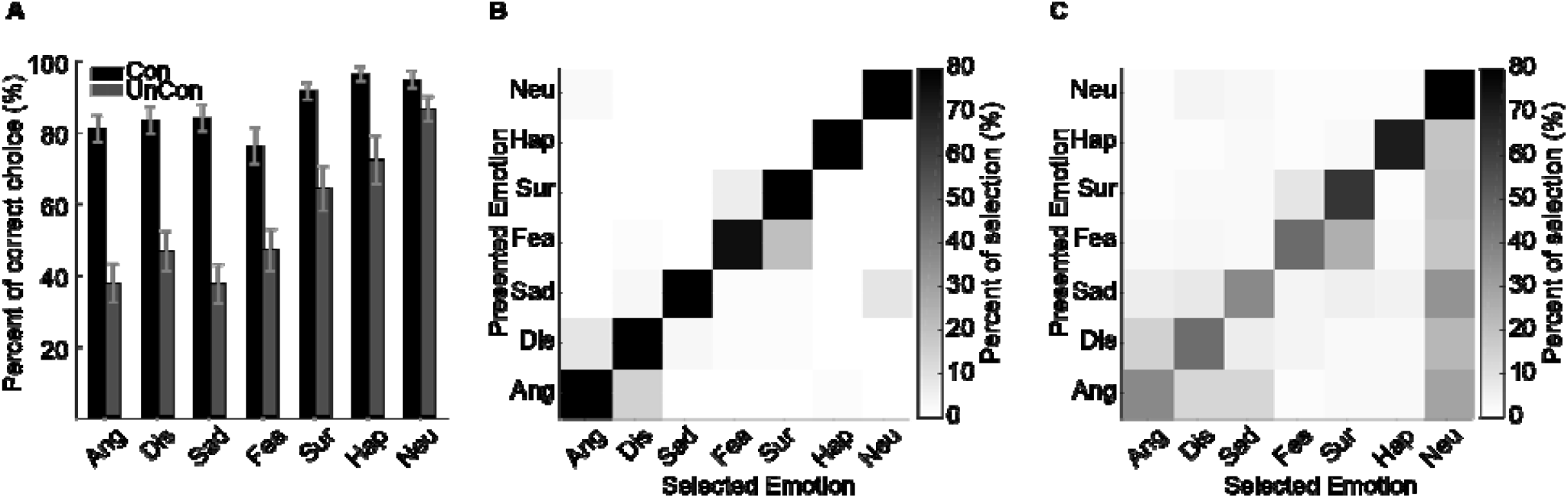
Behavioral performance and confusion matrix. **(A)** The average performance of participants in conscious and unconscious perception for each emotion. Error bars represent the standard error of the mean. As expected, in unconscious perception, performance is lower due to the increased difficulty of the task. **(B)** Confusion matrix for the percentage of selection of emotions in the conscious state. In this figure, the vertical axis indicates the emotion displayed. In contrast, the horizontal axis represents the emotion selected by the participant, and the colours from white to black indicate the selection percentage. The main diagonal determines the correct performance, where the emotions presented and selected are the same. **(C)** Confusion matrix for the percentage of selection of different emotions in the unconscious state. As seen in the figure, there is a significant percentage of neutral responses instead of the displayed emotional expression in all emotions due to the presentation of a neutral mask.

Confusion matrices illustrated performance errors, showing biases towards specific emotions (Figure 2B and C). Challenges in distinguishing fear and surprise (19.95%) and anger and hate (14.58%) were evident in the conscious state matrix (Figure 2B). Notably, misattributions of hate errors to anger (8.59%) and surprise errors to fear (5.87%) occurred. Despite high accuracy in identifying sadness (82.61%) and happiness trials (94.79%), biases were still present. Similar patterns were observed in the unconscious state (Figure 2C), indicating consistent biases between fear and surprise, and anger and hate. Given the neutrality of the mask, participants exhibited discernible errors by selecting the neutral state instead of the correct emotion. Comparisons between the conscious and unconscious states reveal subtle distinctions. For instance, misattribution between sadness and anger, as well as sadness and hate, was notably elevated (5.97 and 7.75, respectively). Correct selection of happiness occurred significantly less frequently than in the conscious state (71.03). Similarly, fear was chosen correctly at a diminished rate (46.02), with 24.72% of the remaining inaccuracies pertaining to surprise.

### Pupillometry Results

Pupil size differences between conscious and unconscious states revealed a pattern of suppression in the conscious state and enhancement in the unconscious state (Figure 3). An initial pupil size increase within the first 500 ms was observed in both states, with suppression attributed to cortical inhibitory mechanisms in conscious trials. Conversely, unconscious trials exhibited an escalating slope in pupil size without the suppression observed in conscious trials.

**FIGURE 3.**
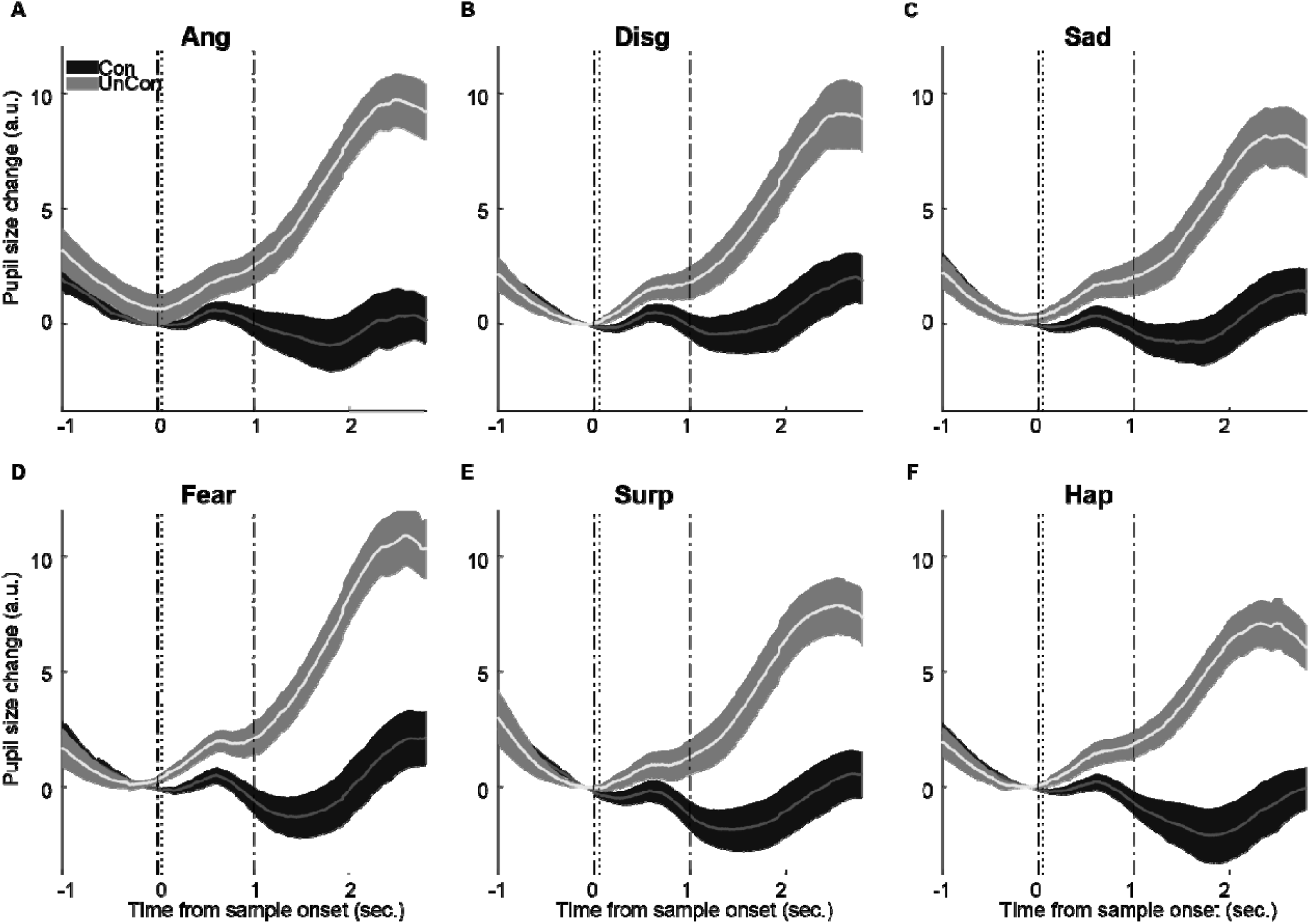
Modulation of pupil size changes for different emotions during conscious and unconscious perception. The lighter shades represent pupil size changes during unconscious detection tasks, while the darker shades represent those during conscious detection tasks. As shown in the figure, all emotions elicit more significant changes in pupil size during unconscious perception than conscious perception. Additionally, there appears to be a difference in pupil dilation for different emotions, and the rise time for each emotion varies. The shaded area in the first plot represents the time interval during which the most significant changes in pupil size occur (between 2000 and 2800 ms after stimulus onset), and it has been used for further analysis.

The peak pupil size for correct trials in both conscious and unconscious states was evaluated in different conditions (Figure 4A). A Wilcoxon signed-rank test revealed significant differences for both states of consciousness for each emotion (p < 0.00), except for neutral (p = 0.07). ANOVA analysis did not indicate significant differences between emotions in the conscious state. However, there was a significant disparity in peak pupil size in the unconscious state. The post-hoc analysis identified this difference specifically between neutral and fear conditions. We evaluated the peak differences in pupil size more concisely using the confusion matrix (Figure 4B-C). The vertical axis denotes the presented emotion, while the horizontal axis represents the selected emotions. The main diameter of the confusion matrices signifies the correct trials.

**FIGURE 4.**
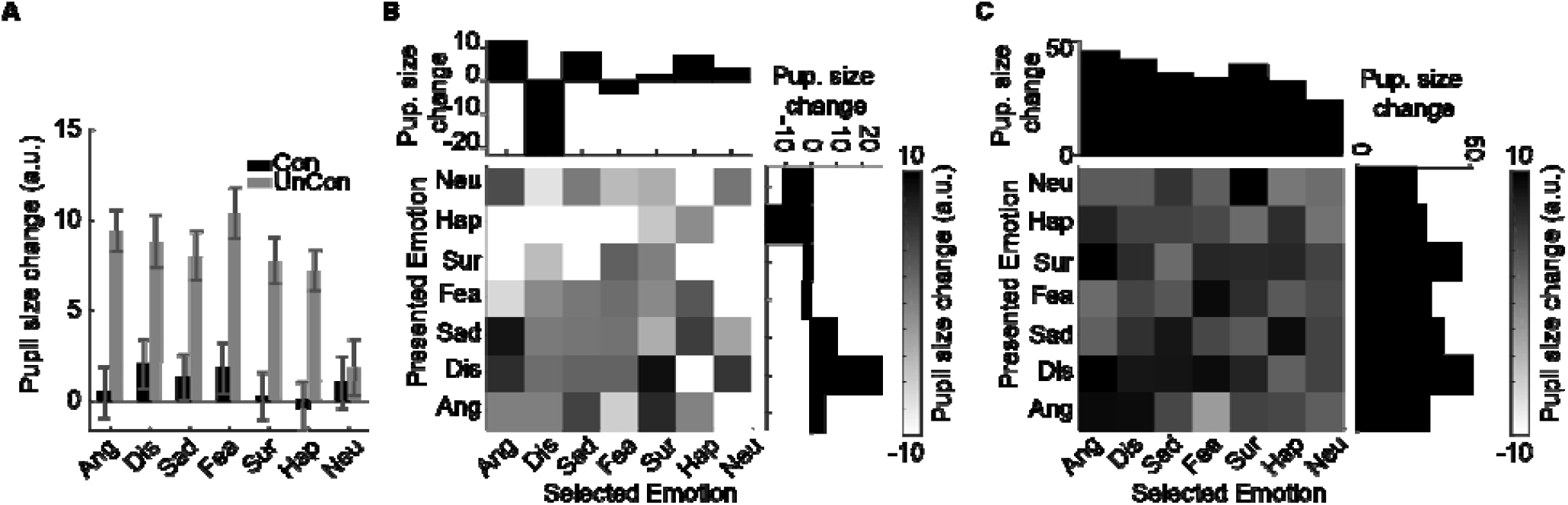
Difference between pupil size changes in different emotions. **(A)** This panel demonstrates the variations in pupil size for trials where the correct answer was selected for both conscious and unconscious tasks. There is a minimal observable change in pupil size for neutral stimuli in both conscious and unconscious states and for emotional stimuli in the conscious state. Error bars represent the standard error of the mean. **(B)** Confusion matrix of pupil size change for different emotions in the conscious state. The upper barplots demonstrate changes in pupil size when the target emotion is selected, irrespective of the particular emotion displayed. The barplots on the right illustrate variations in pupil size when a specific emotion is displayed, regardless of which emotion is selected. **(C)** Confusion matrix of pupil size change for different emotions in the unconscious state.

In the conscious state, when the presented stimulus is positive and the subject mistakenly selects a negative emotion, there is a decrease in pupil size. Also, there are different pupil size changes with the emotion of disgust. In the unconscious state, there is a more significant increase in pupil size overall. The unconscious state exhibits significant pupil size changes, even when emotions are incorrectly selected. Notably, when fear and surprise are chosen correctly or incorrectly, there is a discernible increase in pupil size, with more pronounced changes during negative emotions. Disgust evokes a greater increase when it is the presented emotion. Using peak response alone may not decode the types of emotion and their processing speed accurately. We utilized modeling tools to investigate these parameters more precisely.

### General linear modeling of pupil response

Using the modeling, we investigated the different components of the task in the production of pupil response. Based on the work of Hoeks and Levelt (1993), pupil response models typically assume two points (Hoeks & Levelt, 1993). First, the models assume a stereotyped pupil response function (PuRF) like a gamma function, which describes the time series of pupil dilation in response to a brief event. Second, the models assume that pupil responses to different trial events sum linearly to generate the pupil size time series; that is, they are general linear models (GLMs) (Figure 5). Using this model, we aimed to separate different components and assess their contributions to pupil response. Generally, three important components should be evaluated: task-related response, amplitude, and latency of emotional stimuli presentation.

**FIGURE 5.**
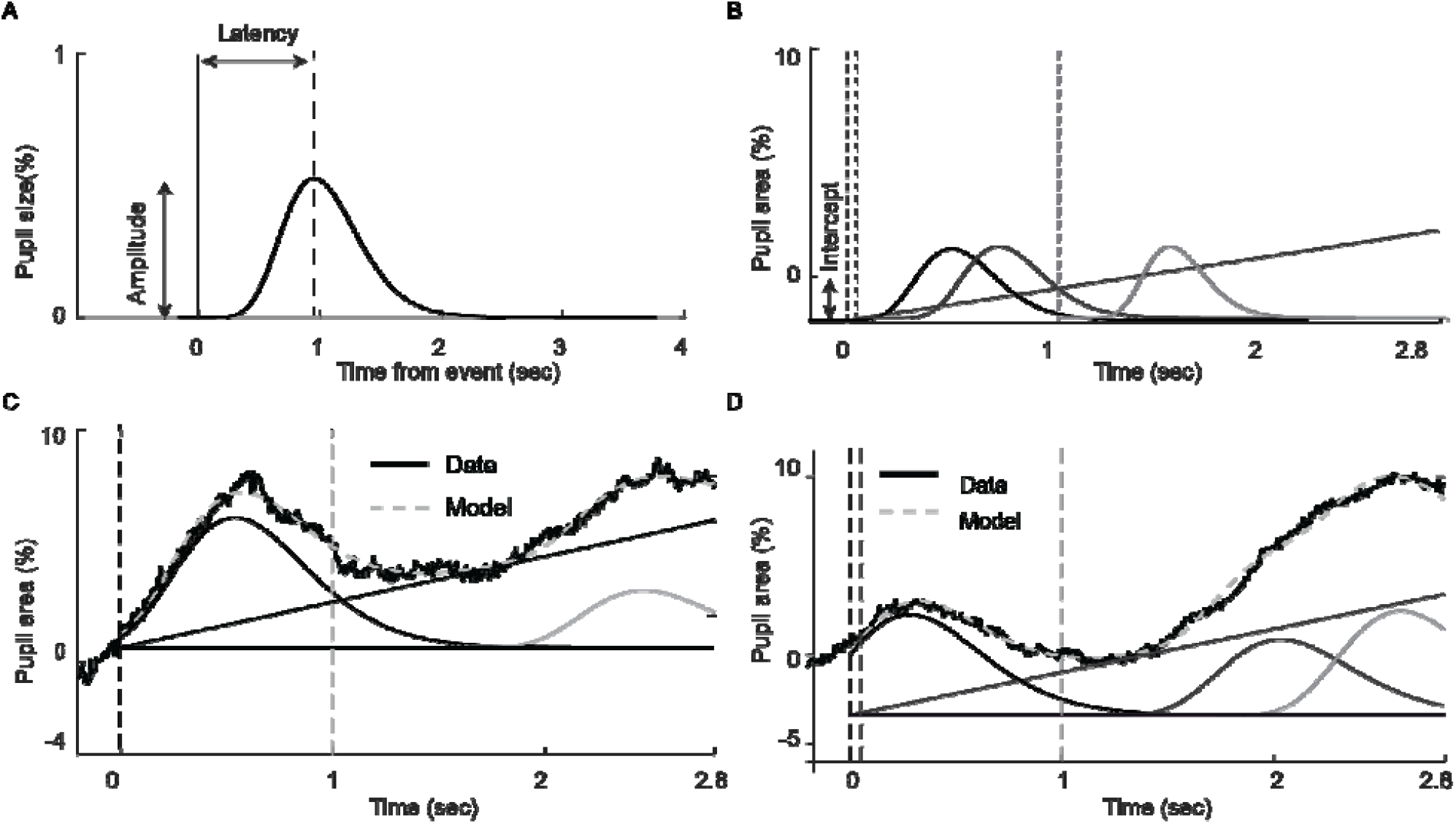
The modeling of the pupil time series. **(A)** a Pupil response function, which describes the pupillary response to an event. The canonical function, a gamma function with n = 10.1 and tmax = 930 ms (vertical dashed line), is shown. **(B)** Trial events hypothesized to drive pupil dilation. Each trial event (visual stimuli) is modeled as a delta function (vertical dashed lines) with some amplitude and some latency with respect to the event. For conscious we have two events at 0 sec, onset of stimulus and 1 sec, offset of stimulus and for unconscious we have one event more presentation of mask at 5o ms after stimulus onset. As line (slop and y-intercept), task-related signal could also be modeled. Shown here is in a solid line. **(C, D)** The pupil time series across (black line) is modeled in two steps. First, each internal signal time series is convolved with the pupil response function to form component pupil responses. Second, the component pupil responses are summed to calculate the model prediction (gray dashed line). Parameters of the model, such as the amplitudes and latencies of the internal event signals, are fit using an optimization procedure. (C) is for sample conscious and (D) is for unconscious trials.

We assumed a linear function as the task-related component for each trial and one baseline shift parameter. These parameters differed for conscious and unconscious states (Figure 6A-B). For the conscious state, angry and surprise have negative slopes, while fear has a positive slope. For the unconscious state, the slope parameter is positive except for neutral, in which there was no emotion. There were no differences in task-related slopes among unconscious emotions. The baseline value is also negative for unconscious emotions and disgust emotion in the conscious state.

**FIGURE 6.**
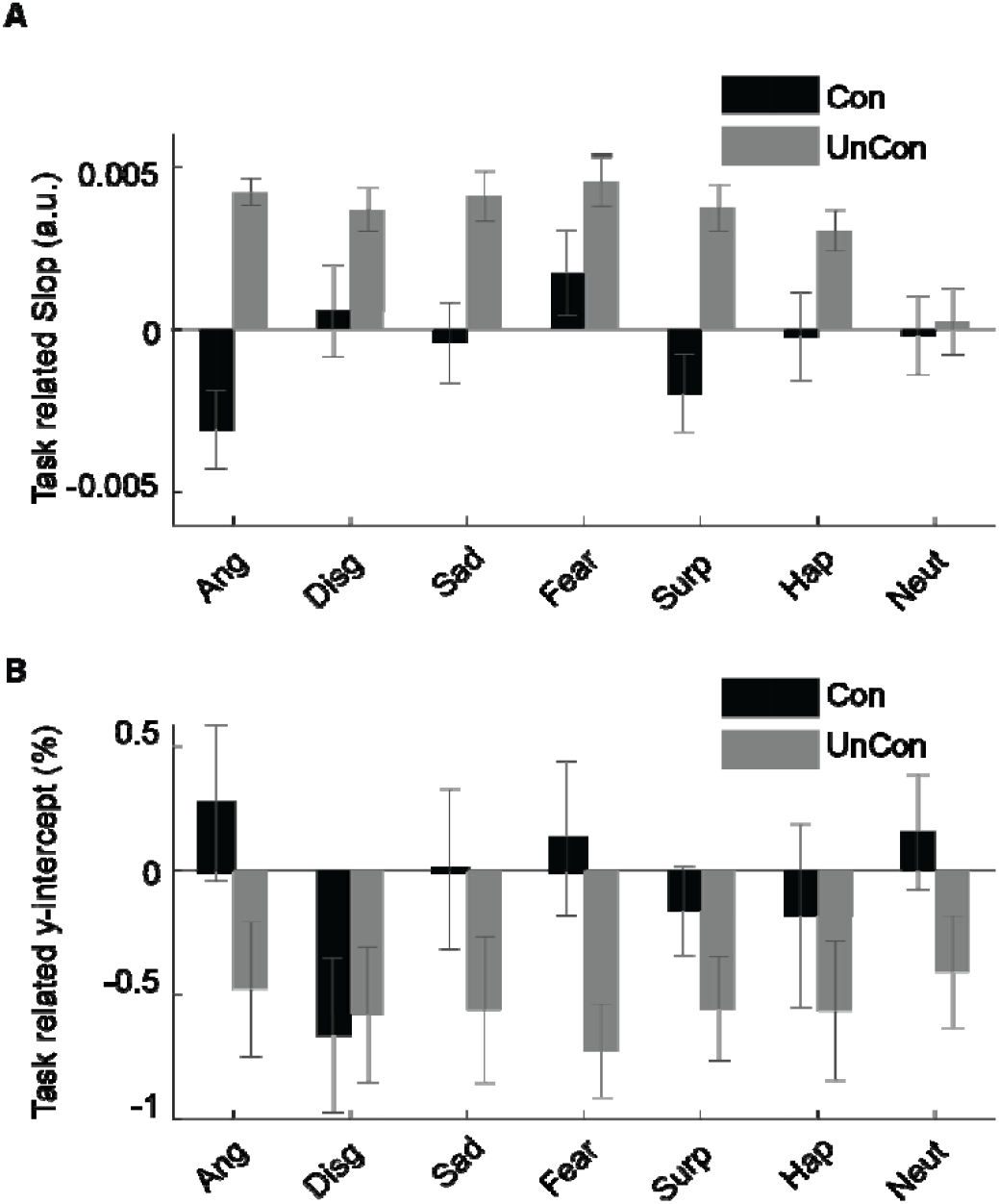
Task related Model parameters over subjects. Tasl related parameters, upper plot slope of line and lower plot y-intercept.

Each event had a response function with an amplitude parameter and a latency parameter associated with it. The amplitude parameter was the value of the nonzero point of the delta function and indicated the magnitude of the internal signal, determining the magnitude of the component pupil response associated with it (Figure 7). The first event for each state is the emotion-related event (amp1 Con and amp1 UuCon). For conscious states, the emotion-related amplitude did not differ between emotions. Similarly, in unconscious states, the emotional response did not differ between emotions.

**FIGURE 7.**
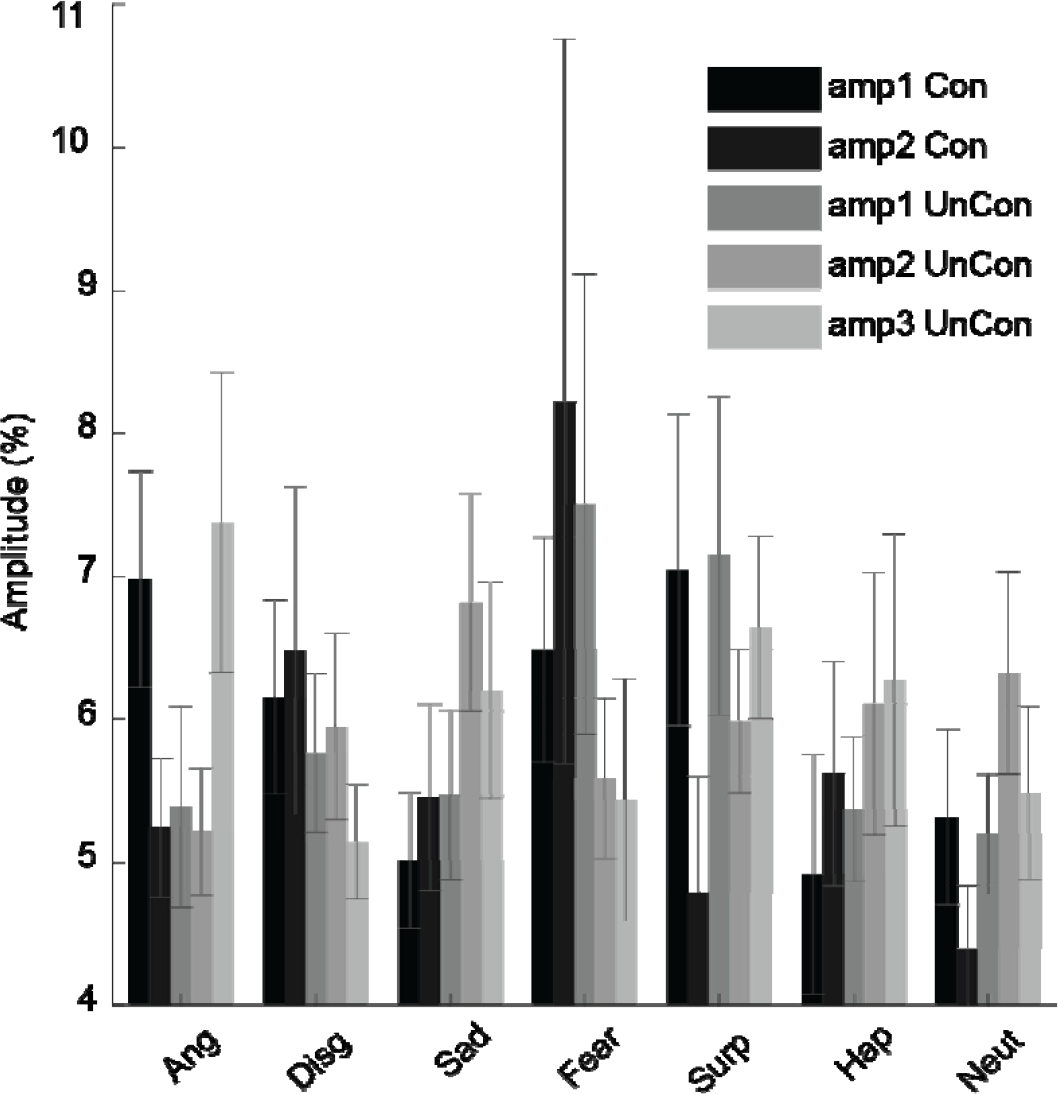
The effect of different events from Model parameters over subjects. Amplitude for different events “amp1 Con” onset stimulus, “amp2 Con” offset stimulus, “amp1 UnCon” onset stimulus, “amp2 UnCon” onset mask and “amp3 UnCon” offset stimulus.

The latency parameter was the time (in ms) of the nonzero value, relative to the time of its corresponding event. The pupil latency reflected the speed of the emotional response (Figure 8). In conscious states, the emotion-related latency did not differ between emotions. Happy and angry were the fastest, while surprise and neutral were the slowest in conscious states. In unconscious states, sad, fear, surprise, and happy were the slowest responses, while disgust was the fastest.

**FIGURE 8.**
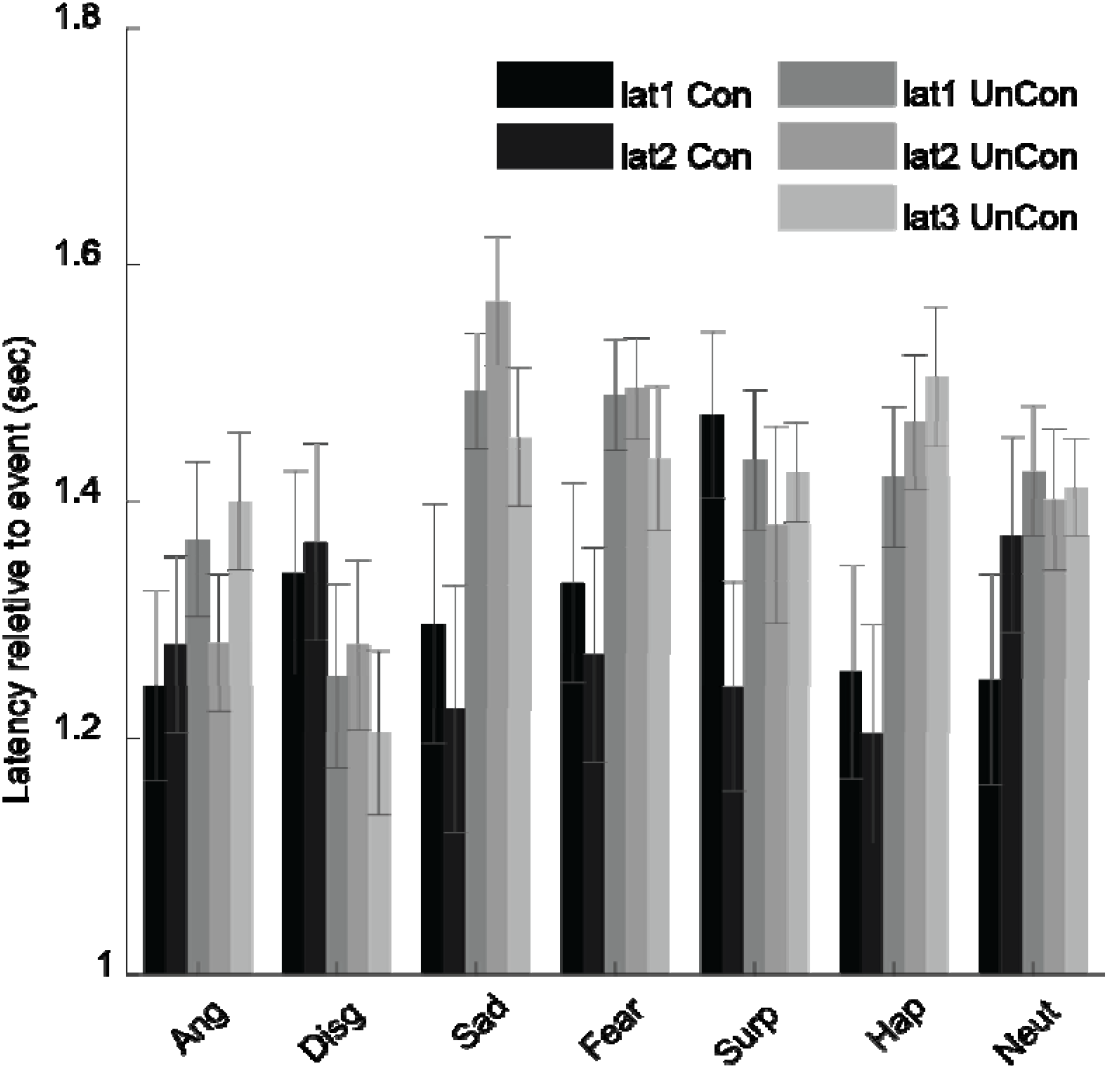
The speed of different events from Model parameters over subjects. latency for different events lat1 Con” onset stimulus, “lat2 Con” offset stimulus, “lat1 UnCon” onset stimulus, “lat2 UnCon” onset mask and “lat3 UnCon” offset stimulus.

## Discussion

The initial noteworthy finding concerning the impact of conscious emotional perception on pupil changes reveals a consistent increase in pupil size during the perception of all emotions, regardless of their valence, in comparison to neutral stimuli. This finding aligns with prior research trends and indicates that emotional perception induces a cognitive load that manifests through observable changes in pupil size. Additionally, Partala et al. (2000) demonstrated that emotional arousal, even in non-visual perceptual contexts such as emotionally engaging sounds, leads to an increase in pupil size (Partala et al., 2000), further supporting the cognitive processing load associated with emotional stimuli. Notably, these pupillary changes are not influenced by the physical properties of the stimuli, as all images were matched in terms of luminance and size.

Transitioning to the unconscious task, the results validate existing research by illustrating that individuals remain sensitive to and affected by other people’s facial expressions even in the absence of conscious awareness (Eastwood & Smilek, 2005). The observed increase in pupil diameter during the response period, compared to the stabilization interval, underscores the influence of emotional states on the autonomic nervous system. Furthermore, a greater magnitude of pupillary changes was noted during unconscious presentations, consistent with prior findings (Tamietto et al., 2009), indicating that unconscious perception elicits heightened arousal when compared to conscious perception, as evidenced by observable changes in pupil size.

Upon closer examination of pupil sizes (Figure 3), a distinct trend emerges between conscious and unconscious trials. Notably, unconscious emotional faces evoke more pronounced changes in pupil size, with a slight decrease observed around 1 second post-onset of unconscious trials. This observation aligns with the proposed hypothesis by Tamietto and de Gelder (2010), suggesting an intricate interplay between cortical and subcortical brain structures during the perception of unconscious emotions (Tamietto & de Gelder, 2010). The hypothesized interaction involves rapid excitatory responses from subcortical structures activating cortical regions, with subsequent inhibitory feedback from the cortex to subcortical areas, accounting for the initial drop in pupil size mediated by the locus coeruleus. Consequently, the paradoxical increase in pupil size during unconscious trials relative to conscious trials may be attributed to the fading inhibitory effect and subsequent activation of excitatory neurons within subcortical structures in response to unconscious emotional stimuli.

It is essential to exercise caution when generalizing results to the broader population, as thresholds and pupil size responses may vary across different demographic groups. Calder et al. (2003) highlighted age-related differences in facial expression recognition, noting that younger individuals generally excel in this domain, while older adults exhibit an advantage in recognizing disgust (Calder et al., 2003). Therefore, the present findings are best interpreted within the context of individuals within a specific age range.

By employing a modeling approach, this study delved into the complex dynamics of pupil sizes, revealing a consistent pattern across conscious and unconscious states and highlighting temporal disparities among emotions. The absence of significant differences in the timing of emotional responses underscores the robustness of the observed patterns and contributes valuable insights into the nuanced interplay between emotional perception and pupil dynamics.

In summary, the discussion underscores the multifaceted nature of emotional perception, elucidates the intricate cortical-subcortical interactions underlying pupil size changes, and emphasizes the importance of considering demographic factors in interpreting pupillary responses to emotional stimuli.

